# Epithelial-mesenchymal transition sensitizes breast cancer cells to cell death via the fungus-derived sesterterpenoid ophiobolin A

**DOI:** 10.1101/2020.05.06.079343

**Authors:** Keighley N. Reisenauer, Yongfeng Tao, Shuxuan Song, Saawan D. Patel, Alec Ingros, Peter Sheesley, Marco Masi, Angela Boari, Antonio Evidente, Alexander V. Kornienko, Daniel Romo, Joseph Taube

**Affiliations:** Department of Biology, Baylor University, Waco, TX, USA; Department of Chemistry and Biochemistry, Baylor University, Waco, TX, USA; Department of Chemical Sciences, University of Naples Federico II, Complesso Universitario Monte Sant’Angelo, Naples, Italy; Institute of Sciences and Food Production, CNR, Bari, Italy; Department of Chemistry and Biochemistry, Texas State University, San Marcos, TX, USA

## Abstract

The epithelial-mesenchymal transition (EMT) imparts properties of cancer stem-like cells, including resistance to frequently used chemotherapy, necessitating the identification of molecules that induce cell death specifically in stem-like cells with EMT properties. Herein, we demonstrate that breast cancer cells enriched for EMT features are more sensitive to cytotoxicity induced by ophiobolin A (OpA), a sesterterpenoid natural product. Using a model of experimentally induced EMT in human mammary epithelial (HMLE) cells, we show that EMT is both necessary and sufficient for OpA sensitivity. Moreover, prolonged, sub-cytotoxic exposure to OpA is sufficient to reduce migration, sphere formation, and resistance to doxorubicin. OpA is well-tolerated in mice and treatment with OpA alone reduces tumor burden. These data identify a driver of EMT-driven cytotoxicity with significant potential for use either in combination with standard chemotherapy or for tumors enriched for EMT features.

## Introduction

Breast cancer patients who have triple-negative breast cancer (TNBC) face poor prognoses driven by high rates of metastasis and early recurrence ^1-6^. TNBC is characterized as histologically negative for estrogen receptor (ER), progesterone receptor (PR), and amplified human epidermal growth factor receptor-2 (HER2), preventing the use of hormone- or HER2-targeted therapies. Instead, treatment with anthracyclinedoxorubicin) and/or taxanes is only capable of providing 5-year survival in about half of TNBC patients ^7-10^.

TNBC is comprised of mostly basal-like and claudin-low intrinsic subtypes, both of which have been characterized as enriched with cancer stem-like cells ^11-13^. Cancer stem-like cells (CSCs) are defined by their ability to re-initiate tumor growth upon transplantation and are hypothesized to fuel metastasis and primary tumor recurrence, resulting in an overall decrease in survival ^14-17^. To improve TNBC patient outcomes, novel and specific approaches targeted at CSCs are needed.

One proposed mechanism driving the emergence of CSC-like cells is the epithelial-mesenchymal transition, EMT ^18,19^. EMT is a trans-differentiation process characterized by spindle-like morphology, loss of apical-basal polarity, increased motility, and a tolerance to anoikis. These phenotypic shifts are driven by gene expression changes mediated by transcription factors Snail (*SNAI1*), Twist (*TWIST1*), and ZEB1, effects of which include upregulation of vimentin and N-cadherin, and downregulation of epithelial markers E-cadherin and miR-200c ^20-26^.

Cells that have undergone an EMT typically acquire CSC properties including decreased sensitivity to conventional chemotherapies used to treat TNBC. This chemoresistance is driven by drug efflux pumps, enhanced DNA repair capacity, mesenchymal-like properties, and epigenetic changes ^16,27-32^. There are currently no approved therapies that specifically target CSCs. A leading pre-clinical compound is salinomycin, reported to decrease the sub-population of CSCs, tumor initiating capability, and chemoresistance, with negligible side effects ^33^. Other naturally occurring compounds such as curcumin, and quercetin have been reported to reduce the effects of EMT by inhibiting key proteins associated with migration (Snail, MMP-2/9), anoikis tolerance (Bcl-2), cell-to-cell adhesion (N-cadherin), and signaling cascades (JAK/STAT, ERK) ^34-37^.

Ophiobolin A (OpA) is a natural product produced from fungi in the genera *Aspergillus, Bipolaris, Cephalosporium, Cochliobolus*, and *Drechslera* ^38^. This sesterterpenoid (25-carbons) is a secondary metabolite that has long been studied for its phytotoxic effects in a variety of plants and has begun to be evaluated as a cytotoxic compound ^38^. Recently published cell culture-based experiments describe a role for OpA in motility inhibition ^39^, membrane depolarization ^40-43^, roles in inflammation ^44^, and reduction in stemness ^45^. *In vivo* data demonstrate that OpA is tolerated in mice and is effective against an orthotopic model of glioblastoma ^40,46,47^. Herein, we investigated the applicability of OpA on EMT-enriched breast cancer and found that experimentally induced EMT enhances the susceptibility of mammary epithelial cells to OpA-induced cell death. Furthermore, breast cancer cell lines treated with OpA experience loss of both stemness and migratory attributes, demonstrating that OpA induces selective cytotoxicity in cells that have undergone EMT. Additionally, OpA is effective in reducing tumor burden in mice with orthotopic, EMT-positive, mammary tumors, highlighting the potential of EMT-targeted cancer treatment.

## Results

### Mammary epithelial cells that have undergone EMT are more sensitive to OpA

Given the previously published link between OpA and CSC-targeted activity ^45^, we investigated a potential link between OpA (Fig. 1A) and EMT using an experimental model of EMT induction. Immortalized human mammary epithelial (HMLE) cells have an epithelial morphology and express E-cadherin. We used HMLE cells, as well as HMLE cells transformed with the Ras oncoprotein (HMLER) that are induced to undergo EMT through lentiviral transduction of viruses driving expression of the EMT-inducing transcript factor Twist, ^26,48^ resulting in the acquisition of a mesenchymal morphology (Fig 1A) and protein expression (Fig. 1B). We measured the Twist-induced selective sensitivity to molecules shown to inhibit CSC properties including salinomycin^49^, ophiobolin A^45^, curcumin^50^, genistein^51^, and disulfiram^52^. Only two such molecules demonstrated selectivity towards EMT-positive cells, salinomycin and ophiobolin A, and only OpA also demonstrated sub-micromolar cytotoxic activity (Fig. 1C). Furthermore, induction of EMT through expression of Twist or through another EMT-TF, Snail 53, in either HMLE or HMLER cells increased sensitivity to OpA-driven cytotoxicity (Fig. 1D). Indeed, the EMT decreased the IC_50_ value from a mean of 137−147 nM for epithelial cells to a mean of 85−91 nM for mesenchymal cells (Fig. 1E). These results stand in stark contrast to EMT-driven resistance to many commonly used chemotherapeutic drugs including doxorubicin, paclitaxel ^49^, and staurosporine (Fig. 1F).

**Figure 1:**
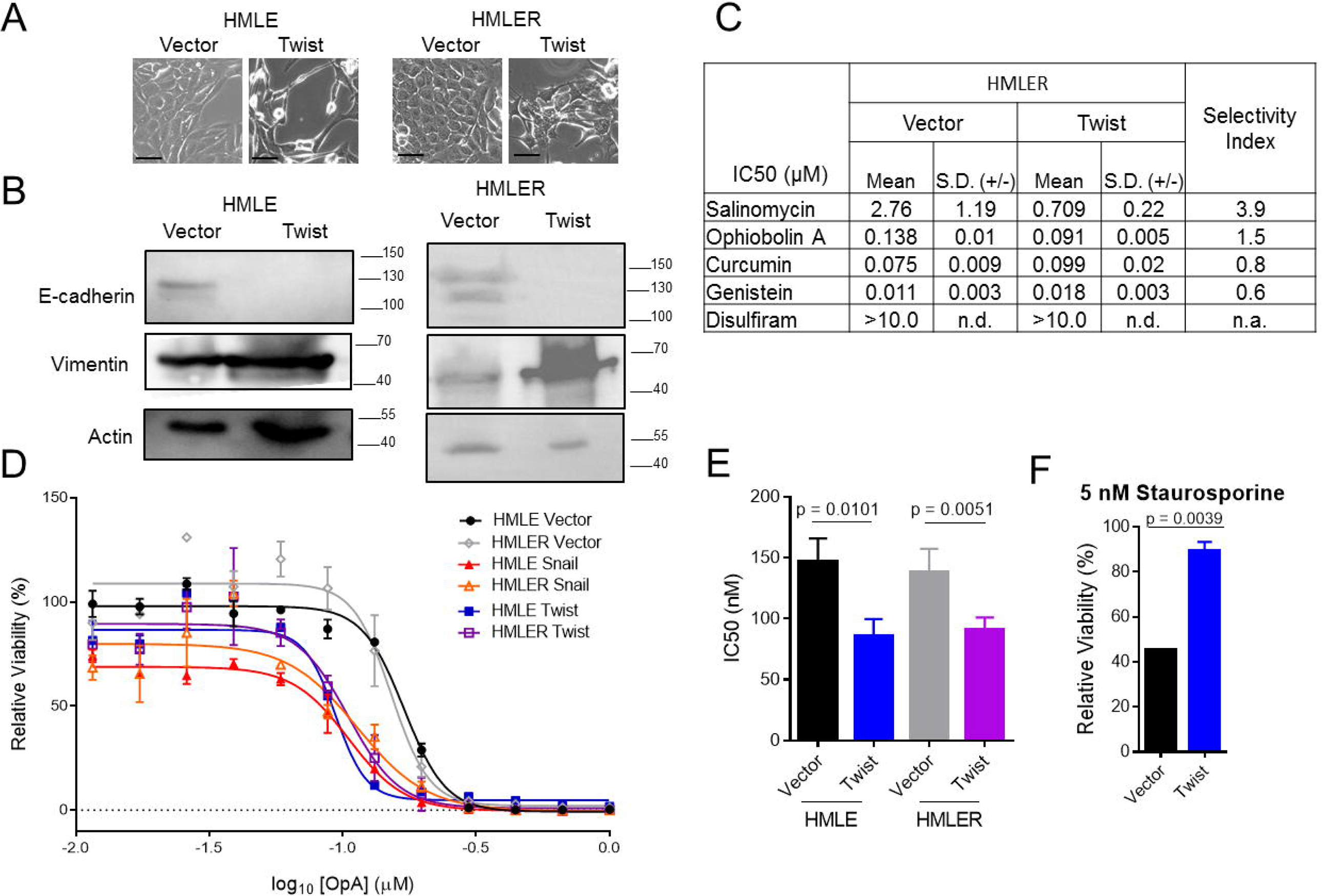
Sensitivity to OpA is enhanced by EMT. (A) Representative morphology of non-transformed, immortalized, mammary epithelial cells expressing Twist or control vector. Scale bar represents 20 µm. (B) Representative western blot showing E-cadherin and vimentin protein expression. Images cropped to show relevant bands. Full-length blots available in supplemental information. (C) Cytotoxic activity of the indicated compounds was measured, in triplicate, by MTS assay. Mean and standard deviation of IC50 values are reported. Selectivity index is calculated as (HMLE Vector IC50) / (HMLE Twist IC50). (D) Representative data indicating cytotoxicity as for the indicated cell lines. Error bars represent standard deviation. (E/F) Mean and standard deviation of IC50 values for OpA (E), n = 3 or 4, unpaired t-test and staurosporine (F), n = 2, unpaired t-test. n.d. = not determined, n.a. = not applicable.

### miR-200c suppression is necessary for sensitivity to OpA

Because we observed that OpA selectively impacts cells that have undergone EMT, we next evaluated whether reversing the EMT status of these cells would be sufficient to restore OpA sensitivity. To do this, we introduced a ZEB1-targeting microRNA into HMLE-Twist and HMLER-Twist cells. miR-200c expression has been shown to be sufficient to reverse EMT ^54^. We verified over-expression of miR-200c driven by transduction with a lentiviral vector (Fig. 2A). We next measured sensitivity to OpA and found that induction of miR-200c partially restored resistance to OpA (Fig 2B). This indicates that expression modulation of the EMT state impacts sensitivity to OpA.

**Figure 2:**
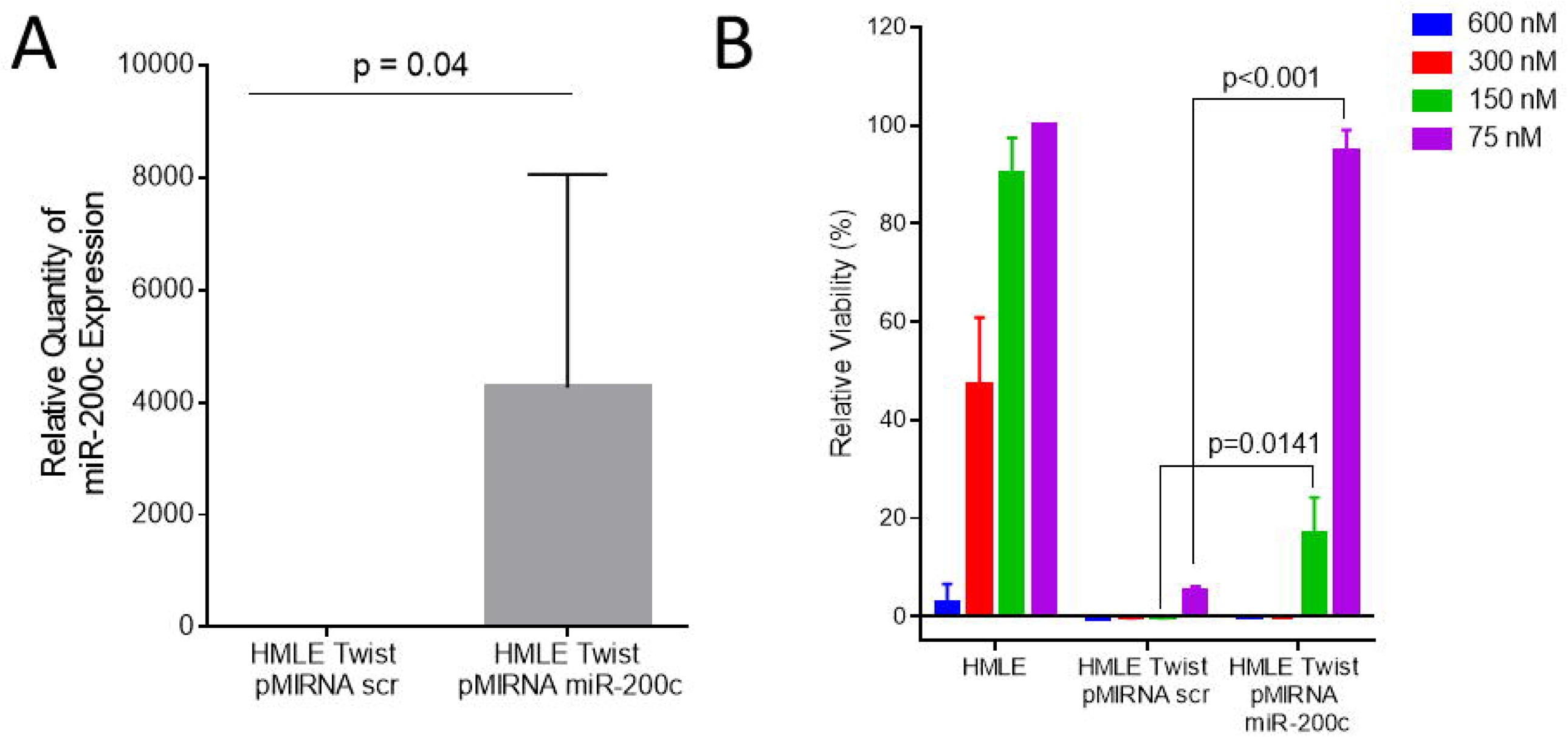
The EMT state is necessary for sensitivity to Ophiobolin. (A) Mean and standard deviation of miR-200c expression in HMLE Twist cells expressing ectopic miR-200c or a control vector. N=3 (B) Mean and standard deviation of relative viability for indicated doses of OpA in HMLE and HMLE Twist cells expressing ectopic miR-200c or a control vector. n=3, unpaired t-test.

### Persistent treatment with OpA alters cellular phenotypes

Engaging the EMT program can confer stemness properties in cancer cells (Mani et al., 2007; Morel et al., 2008). To examine the effect of OpA on breast cancer cells, we measured the cytotoxic activity on the ER-positive, CSC-poor, epithelial-like MCF7 and triple-negative, CSC-rich, mesenchymal-like MDA-MB-231 cell lines. While both cell lines were highly responsive to OpA, the MDA-MB-231 cells displayed significantly greater cell death at an 80 nM dose (Fig. 3A). To evaluate the impact of sub-cytotoxic doses of OpA on EMT and CSC phenotypes, we performed experiments on CSC-rich MDA-MB-231 cells using continuous, multi-day treatment of 30 nM OpA or 30 nM deoxy-OpA, as a negative control (Fig. 3B-black arrow). Continuous treatment with a sub-cytotoxic doses of OpA, but not 3-deoxy OpA triggered modest changes in cell morphology toward a more compact and cobblestone-like appearance, characteristic of epithelial cells (Fig. 3C). EMT is necessary for the migratory capacity of MDA-MB-231 cells. To ascertain if OpA inhibited migration we performed a wound healing assay. Consistent with an effect on EMT properties, cells pre-treated with sub-cytotoxic doses of OpA, but not 3-deoxy-OpA, failed to migrate in response to a scratch wound (Fig. 3D,E). We next tested CSC-characteristic anchorage-independent growth using a mammosphere assay. Consistent with an effect on CSC properties, we observed that pre-treatment of MDA-MB-231 cells with OpA reduced sphere formation (Fig 3F). In summary, persistent treatment of a CSC-rich breast cancer cell line with OpA diminishes sphere formation and migratory properties associates with CSC and EMT.

**Figure 3:**
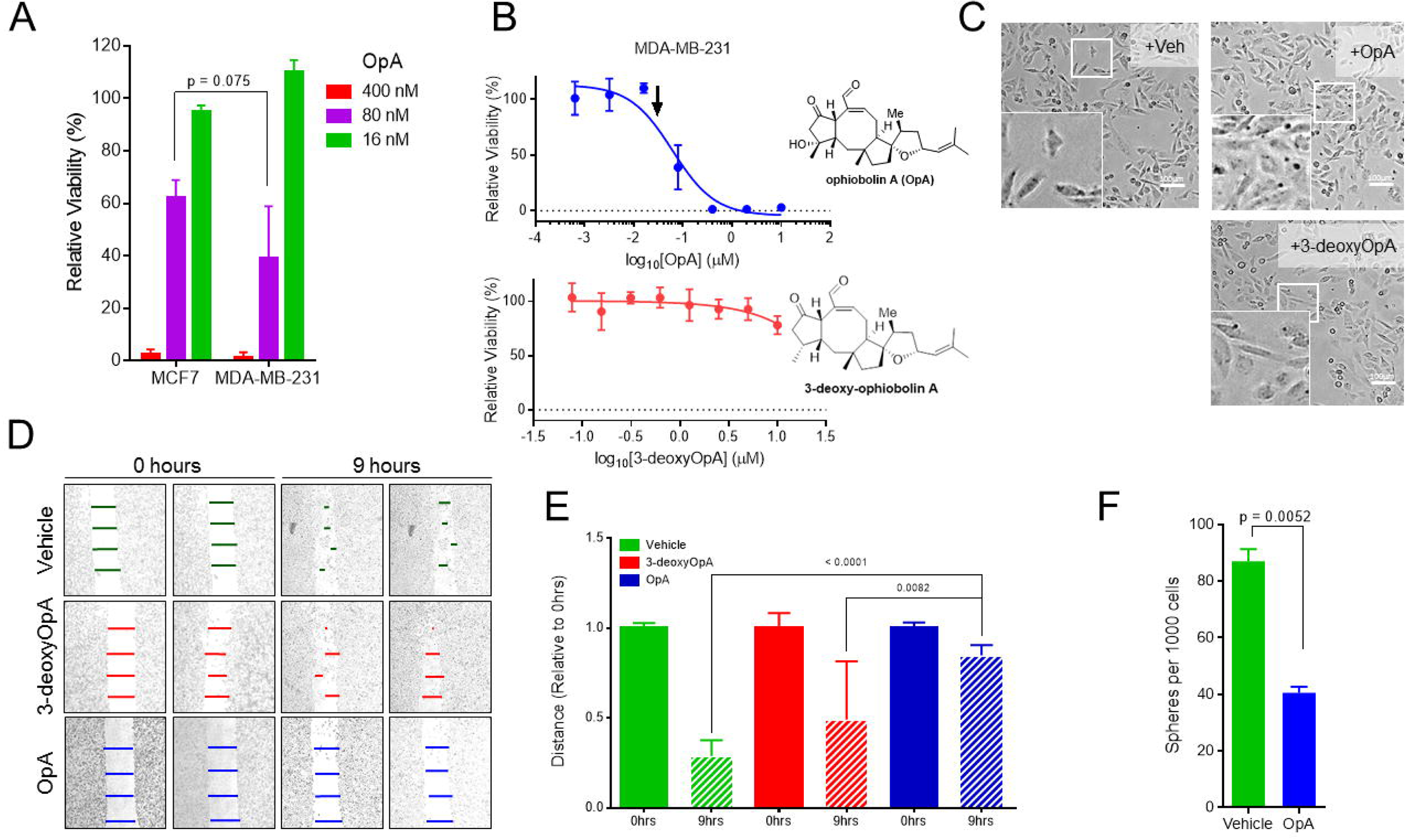
Treatment with OpA, but not an inactive congener, suppresses EMT-driven cell behavior. (A) Mean and standard deviation of relative viability for indicated doses of OpA in MCF7 and MDA-MB-231 cells, n = 8, unpaired t-test. (B) Representative data indicating cytotoxicity of MDA-MB-231 to the indicated compounds. Black arrow indicates dose used for sub-cytotoxic dosing. (C) Representative morphology of MDA-MB-231 cells were treated with the indicated compounds at 30nM for 30 days. Scale bar = 100 µm. (D/E) MDA-MB-231 cells, treated with the indicated compounds at 30nM for 4 days, were cultured in clean media for 12 hours then subjected to a wound healing assay. (F) MDA-MB-231 cells, treated with the indicated compounds at 30nM for 10 days, were cultured in clean media for 48 hours then subjected to a sphere-forming assay. n=8, unpaired t-test

### OpA treatment increases sensitivity to chemotherapy

EMT-promoted stemness drives resistance to commonly used chemotherapies. One approach to overcoming this problem is to consider dual-treatment therapies that combine CSC-targeting compounds with conventional drugs. To examine the combinatorial impact of OpA treatment we co-treated MDA-MB-231 cells with a dilution series of OpA and either doxorubicin or paclitaxel. Co-treatment with as little as 12.5 nM OpA enhanced the cytotoxic response from doxorubicin (Fig. 4A), while 50 nM OpA enhanced the cytotoxic response from paclitaxel (Fig. 4B). Notably, addition of 50 nM OpA was sufficient to maintain cytotoxic activity despite a 25-fold reduction in the dose of doxorubicin (Fig. 4A-orange bar) and a 5-fold reduction in the dose of paclitaxel (Fig. 4B-orange bar). Indeed, when analyzed using Combenefit, these dose combinations tended toward synergistic effects (Fig. 4C,D). Combination treatment using 3-deoxy-OpA did not result in altered activity (Fig. 4E,F) The capacity of OpA to act in concert with clinically useful chemotherapeutic agents indicates that co-treatment may be useful to more effectively treat breast cancer.

**Figure 4:**
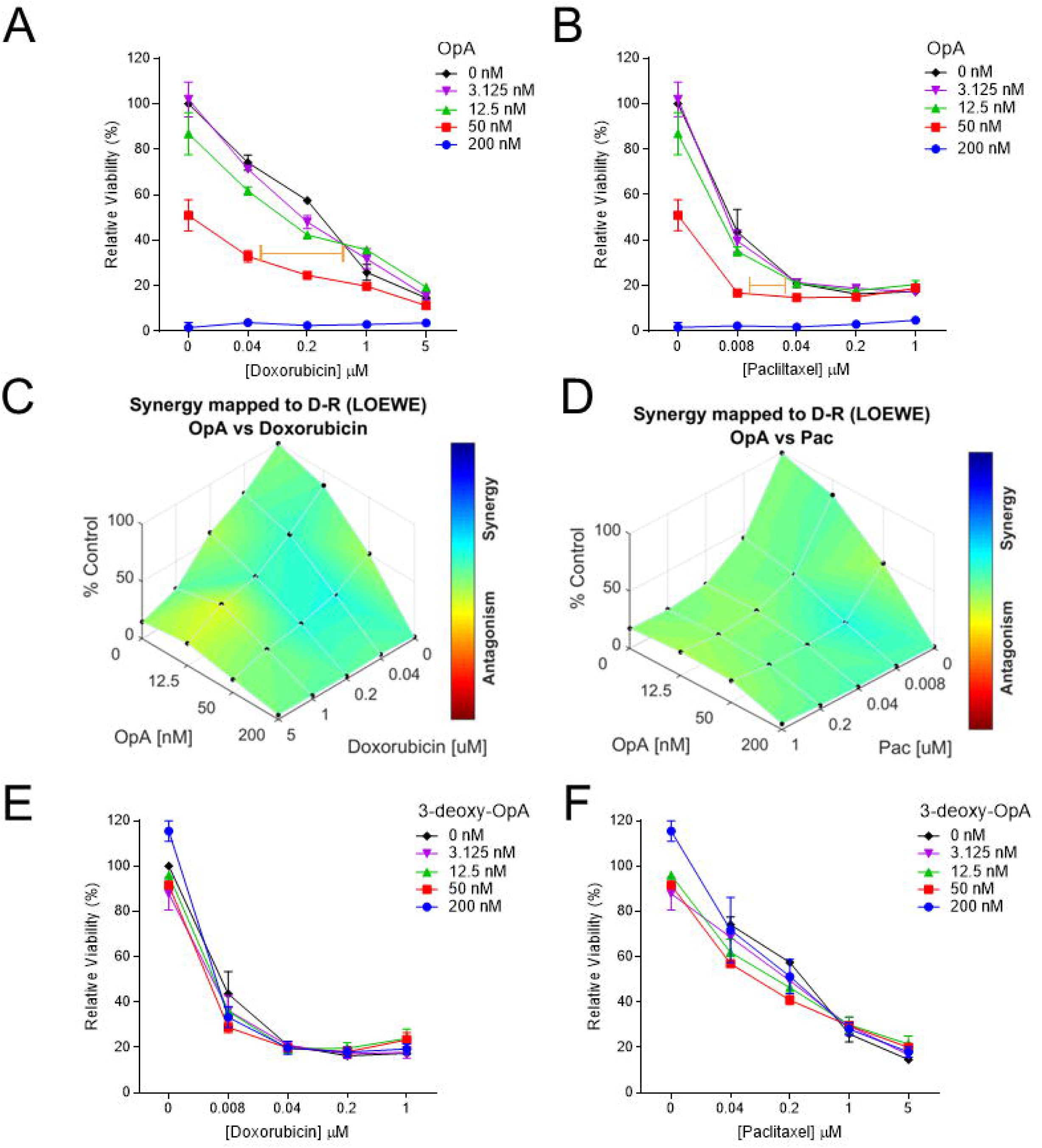
Combinatorial activity for OpA with doxorubicin and paclitaxel. (A/B) Representative data indicating cytotoxicity of MDA-MB-231 to a range of doses for OpA and doxorubicin (A) or OpA and paclitaxel (B). (C/D) Data from (A/B) are represented using Combenefit. Blue-shaded areas represent dose combinations with synergistic effects (E/F) Representative data indicating cytotoxicity of MDA-MB-231 to a range of doses for 3-deoxy-OpA and doxorubicin (E) or 3-deoxy-OpA and paclitaxel (F).

### OpA is tolerated *in vivo* and suppresses Twist-expressing tumor growth

We next assessed whether OpA treatment alone is sufficient to reduce tumor growth in mice. In order to gauge the impact of OpA on tumor growth, immunocompromised mice were orthotopically injected with Ras-transformed HMLE cells expressing the Twist transcription factor (HMLER-Twist) to induce EMT. Following the emergence of palpable tumors, mice were randomly assigned to either the control (DMSO diluted into saline) or OpA treatment groups. Thrice weekly injections for 3 weeks consisting of 10 mg/kg of OpA were not well tolerated as mice exhibited weight loss greater than 20% of initial body weight and two adverse outcomes were recorded prior to the final dose (Fig. 5A). However, a dose of 5 mg/kg was better tolerated with final weight loss less than 15% and one adverse outcome while a dose of 2.5 mg/kg had no statistically significant impact on body weight (Fig. 5A). A dose of 5 mg/kg of OpA was sufficient to significantly reduce the endpoint tumor volume of HMLER-Twist tumors (Fig. 5B).

**Figure 5:**
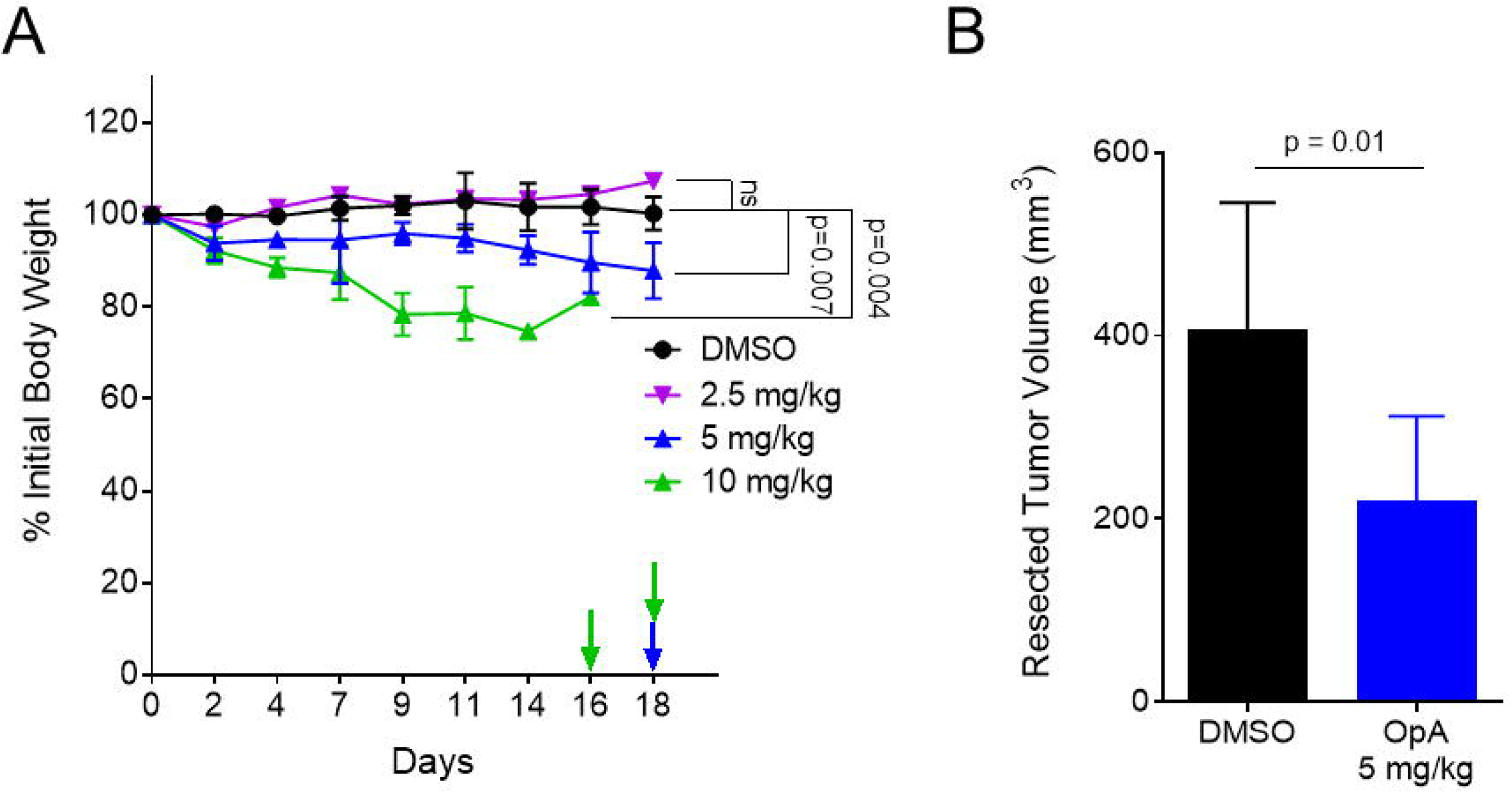
OpA is tolerated in vivo and suppresses tumor growth from cells over-expressing Twist. (A/B) Mice, bearing tumors composed of HMLE-Ras Twist cell, were injected with 10 mg/kg (n=2), 5 mg/kg (n=5), 2.5 mg/kg (n=3), of OpA, or vehicle control (n=8), thrice weekly for three weeks. (A) Body weight was tracked. Arrows indicate endpoint criterion met for an individual animal. Statistical significance measured using the Holm-Sidak method with an alpha of 5%. (B) End-point tumor volume was compared via unpaired t-test, n=4.

## Discussion

Currently, conventional chemotherapeutic drugs are able to elicit high response rates in about half of TNBC patients; however, the remaining patients eventually develop progressive disease^2^, with some even experiencing more aggressive and CSC-rich tumors after therapy^14,55^. Identification of molecules that mediate sensitivity to CSC-rich cell populations will facilitate the development of novel therapies and may improve responses to currently available therapies.

While several other natural products have been linked to CSC-targeting^34,35,49,56-61^, our work highlights a natural product that selectively kills breast CSCs exhibiting EMT features. We demonstrate selective and necessary sensitivity to OpA in cells that have undergone EMT. Further, we show a reduction of EMT phenotypes such a migration and morphology, as well as reduction in sphere-forming capacity in a TNBC cell model. Extending OpA’s efficacy in reducing CSC-related properties, our data suggest increased sensitivity to conventional chemotherapeutics doxorubicin and paclitaxel when co-treated with OpA. Finally, we evaluate the efficacy of OpA *in vivo* and show a high level of tolerance to OpA, as well as volume reduction in an EMT-positive, primary tumor.

Evolution-driven selection of natural products imparts biological activities useful for disease treatment and which may not be mimicked by selective kinase inhibitors. Other successful natural products that have driven cancer therapies include taxol, vinblastin, anthracyclines, daunomycin and doxorubicin^62^. Many studies^40,42,43,47,63-66^ have evaluated one such natural product, OpA, in cancer settings, predominantly using *in vitro* models, and, similar to our present study, these studies report IC_50_ values in the low nanomolar range. Our work is one of the first to evaluate OpA *in vivo* and is the first to describe the impact of EMT on OpA sensitivity. By focusing on the effects on EMT and stemness phenotypes, this work opens the door for the discovery of essential molecular pathways and for the investigation of OpA derivatives as a future cancer treatment.

## Materials and Methods

### Cell Lines

MCF7 were received from ATCC. MDA-MB-231, Hs578T, HMLE, HMLER, HMLE Twist, and HMLER Twist were kindly gifted from Dr. Sendurai Mani (MD Anderson Cancer Center).Breast cancer cells were cultured in Dulbecco’s Modified Eagle’s Medium (DMEM) (Corning Inc., Kennebuck, ME, USA) supplemented with 10% fetal bovine serum (FBS) (Equitech-Bio Inc., Kerrville, Texas, USA) and 1X antibiotics (Penicillin/Streptomycin, Lonza, Basel, Switzerland). Immortalized human mammary epithelial cells (HMLE) and derivatives were maintained as in Elenbaas et. al ^67^. Cell lines were tested monthly for mycoplasma and validated via STR testing. Culture conditions were 37 °C, 5% CO_2_.

### Reagents

OpA was produced by fermentation of the fungus *Drechslera gigantea*. It was extracted from the fungal culture filtrates, purified, crystallized and identified by ^1^H NMR and ESI MS as previously reported^68^. The purity of OpA was >98% as ascertained by ^1^H NMR and HPLC analyses.

3-Deoxy OpA was synthesized from ophiobolin I^69,70^ which was also obtained through fermentation as previously reported^68^. A two-step synthetic sequence involving conjugate reduction of the enone which proceeded with high diastereoselectivity (>19:1 by 600 MHz ^1^H NMR) followed by a Ru(IV)-mediated oxidation of the primary alcohol to the aldehyde delivered 3-deoxy OpA. It should be noted that the methyl group at C3 is epimeric with respect to the C3-methyl group in OpA. However, the importance of the C3-hydroxy group and/or the stereochemistry of this methyl group was verified through studies described below and 3-deoxy OpA served as a negative control. Further details are provided in Supplemental Figure 1.

### Viability

Cells were plated with 2,000 cells per well in a 96-well plate and allowed to adhere overnight. Compounds, suspended in DMSO and diluted into PBS, or vehicle were added to the culture medium and incubated for 72 hours at 37 °C, 5% CO_2_. Following manufacturer suggested protocol, 20 μL CellTiter 96® AQ_ueous_ One Solution Cell Proliferation Assay (MTS; Promega, Madison, WI, USA) was added and incubated 1−4 hours at 37 °C, 5% CO_2_. Absorbance was measured at 490 nm using a 96-well plate reader (Fisher Scientific, Hampton, NH, USA).

### RNA extraction and detection

Cells were lysed in the presence of Trizol® Reagent (Thermo Scientific, Waltham, MA, USA) and total RNA extracted following manufacturer protocol recommendations. Relative quantification of the mRNA levels was performed using the comparative Ct method with Taqman assays for U6 as the reference gene for microRNA analysis and SYBR Green assays GAPDH as the reference gene for mRNA analysis, and with the formula 2^-^ΔΔCt (Applied Biosystems, Foster City, CA, USA; Thermo Scientific). All quantitative reverse transcription-PCR (RT-PCR) experiments were run in triplicate and a mean value was used for the determination of mRNA levels.

### Western blotting and antibodies

Cells were lysed in the presence of 100 microliters radio-immunoprecipitation (RIPA) buffer containing protease inhibitors (Alfa Aesar, Stoughton, MA, USA) on ice. Protein was quantified using the Bradford Assay (BioRad, Hercules, CA, USA). Twenty micrograms of total protein from each sample was resolved on a 4%–12% SDS-PAGE gel and transferred to PVDF membranes. Sister blots were then probed with antibodies including anti-E-cadherin (Cell Signaling, Danvers, MA, USA), anti-Fibronectin (Sigma-Aldrich, St. Louis, MO, USA), anti-vimentin (Protein Technologies, Tucson, AZ, USA), anti-N-cadherin (Cell Signaling), anti-ZEB1 (Santa Cruz Biotechnologies, Dallas, TX, USA), or anti-β-actin (BD Biosciences, San Jose, CA) antibody. Chemiluminescent signals were detected with ECL™ prime (Thermo Fisher Scientific) using the Biorad ChemiDoc system. If necessary, blots were stripped with ECL Stripping Buffer (Li-Cor, Lincoln, NB, USA) following manufacturer protocol. Bands were quantified using ImageJ.

### Mammosphere Assay

Cells were harvested according to standard protocol and were suspended in serum-free mammary epithelial growth medium (MEGM) supplemented with 1% methyl cellulose, 20 ng/mL FGF, 10 ng/mL EGF, and 4 μg/mL heparin. Cells were plated in 4 replicates in a flat-bottom ultra-low attachment 96-well plate (Corning) and allowed to grow at 37 °C, 5% CO_2_ for 10−14 days and were monitored microscopically to ensure that they did not become confluent during the experiment. 100 µL low-attachment media was added after 7 days. Wells were imaged using 4x magnification on a computer-assisted phase contrast microscope (Nikon, Tokyo, Japan). Spheres larger than 100 micrometers were counted.

### Migration

For migration cells were serum-starved overnight and scratch wounds were created using a sterile pipette tip on the cell monolayer or by plating cells in 2-well culture inserts (Ibidi, Madison, WI). Cell migration rates were determined by measuring the distance between cell fronts after the indicated number of days in culture. The distance between the two edges at multiple points was quantified using ImageJ at the indicated timepoints.

### Co-treatment and interaction

Cells were treated with doxorubicin (Selleckchem, Houston, TX, USA), paclitaxel (Selleckchem), OpA, 3-deoxy OpA, or matched-percentage DMSO in decreasing concentrations and incubated for 72 hours before measuring viability using MTS. Interactions were quantified using the Combenefit program with the Loewe model and dose-response surface mapping^71^.

### Tumor growth

Female Scid/bg (CB17.Cg-PrkdcscidLystbg-J/Crl) mice (5–8 weeks old) were obtained from Charles River Laboratories (Wilmington, MA, USA). Animals were maintained under clean room conditions in sterile filter top cages with autoclaved bedding and housed on high efficiency particulate air–filtered ventilated racks. Animals received sterile rodent chow and acidified water *ad libitum*. All of the procedures were conducted in accordance with the Institute for Laboratory Animal Research Guide for the Care and Use of Laboratory Animals and with Baylor University Animal Care and Use Committee guidelines. HMLE-Ras + Twist cells were harvested, pelleted by centrifugation at 2000xg for 2 minutes, and resuspended in sterile serum-free medium supplemented with 30% to 50% Matrigel (BD Biosciences, San Jose, CA, USA). Cells (2×10^6^ in 100 aliquots) were implanted into the left fourth mammary fat of each mouse and allowed to grow to the designated size, as measured by caliper, before the administration of OpA at 5 mg/kg or 10 mg/kg three times weekly for three weeks. Tumor volume and body weight were recorded concurrently to injection protocol ^72^. At designated times, mice were humanely euthanized, and tumors were collected. Experiments were approved by Baylor University IACUC (#1441130).

### Statistical analysis

Unless otherwise stated, statistical differences were determined using a student’s T-test. The GraphPad PRISM software v6 was used to perform these analyses. Statistical significance levels are annotated as n.s. = non-significant, *p < 0.05, **p < 0.01, ***p < 0.001, ****p < 0.0001.

## Acknowledgements

We acknowledge the entire Taube Lab for invaluable discussion and advice. Also, we appreciate the assistance of Dr. Igor Bado from Baylor College of Medicine for advice regarding Combenefit. Graphical abstract created with BioRender. The authors also thank Dr. Maurizio Vurro (Institute of Sciences and Food Production, CNR, Bari, Italy) for the supply of *Drechslera gigantea* culture filtrates. This work was supported by the Cancer Prevention and Research Institute of Texas, grant #RP180771 to J.H.T. and D.R.

## Author Contributions

Experiments performed by K.N.R. with contributions from S.S., P.S., S.P., and A.I. Isolation and characterization of ophiobolins by M.M., A.B., A.E., and A.K. Novel syntheses by Y.T. and D.R. Study design, manuscript drafting by K.N.R. and J.H.T., with editing by D.R. and A.K. All authors read and approved the final manuscript.

