## Supplemental Data for "Epithelial-mesenchymal transition sensitizes breast cancer cells to cell death via the fungus-derived sesterterpenoid ophiobolin A"

**Affiliations**

**Supporting Information**

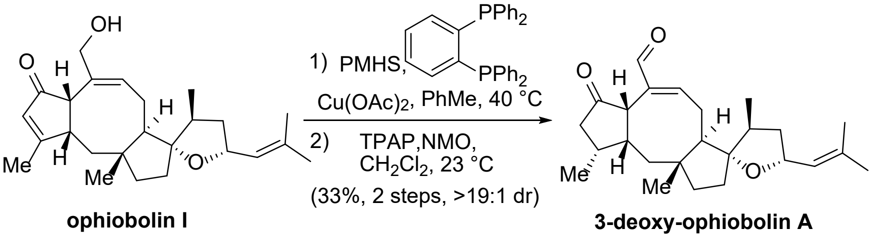


**Synthesis of 3-deoxy-ophiobolin A**. To a dry 1-dram vial was added 3 mL dry PhMe, $Cu(OAc)_2 \bullet H_2O$ (13 mg, 0.065 mmol, 5 equiv), 1,2-bis(diphenylphosphino)benzene (3.0 mg, 0.007 mmol, 0.5 equiv), *t*-BuOH (0.18 mL, 1.95 mmol, 150 equiv), polymethyl hydrosilane (PMHS) (0.12 mL, 1.95 mmol, 150 equiv). The mixture was stirred at 23 °C under argon for 30 min to generate a stock solution of the required reducing agent. In another dry 1-dram vial, ophiobolin I[1] (3.0 mg, 0.008 mmol, 1 equiv) was dissolved in 0.5 mL dry PhMe, and 0.3 mL of the above reagent stock solution was added. The reaction was heated to 40 °C under argon and stirred at this temperature for 2 h. The mixture was concentrated and filtered through a pad of silica gel with $CH_2Cl_2$ as eluent. The filtrate was then concentrated, dissolved in $CH_2Cl_2$, and then *N*-methyl morpholine *N*-oxide (NMO, 1.2 mg, 0.01 mmol, 2 equiv) and *N*-tetra n-propyl ruthenium tetroxide (TPAP, 0.1 mg, 0.0003 mmol, 0.05 equiv) were added and stirred at 23 °C for 1 h. The mixture was concentrated and analysis of the crude $^1$H NMR (600 MHz) suggested formation of a single diastereomer during the conjugate reduction (>19:1). The mixture was then purified by silica gel chromatography (0 → 10% EtOAc/hexanes) to afford 3-deoxy ophiobolin A (1.0 mg, 33%, >19:1 dr by $^1$H NMR) as a colorless residue: TLC (EtOAc: hexanes, 1:2 v/v): $R_f$ = 0.3; $^1$H NMR (600 MHz, $CDCl_3$) δ 9.15 (s, 1H), 6.78 (dd, *J* = 7.0, 2.4 Hz, 1H), 5.06 (dq, *J* = 9.2, 1.5 Hz, 1H), 4.54 (dt, *J* = 8.6, 7.1 Hz, 1H), 3.08 (d, *J* = 9.4 Hz, 1H), 2.72 (dt, *J* = 19.8, 3.1 Hz, 1H), 2.53 (dd, *J* = 14.2, 4.2 Hz, 1H), 2.48 (dd, *J* = 17.3, 12.6 Hz, 1H), 2.36 (ddd, *J* = 17.3, 6.6, 1.6 Hz, 1H), 2.30 – 2.22 (m, 1H), 2.14 (q, *J* = 6.9 Hz, 1H), 1.91 (dd, *J* = 13.6, 3.5 Hz, 1H), 1.76 – 1.67 (m, 7H), 1.64 (d, *J* = 1.4 Hz, 3H), 1.60 (d, *J* = 1.3 Hz, 3H), 1.36 – 1.25 (m, 2H), 1.09 (d, *J* = 5.8 Hz, 3H), 0.97 (d, *J* = 6.9 Hz, 3H), 0.76 (s, 3H); $^{13}$C NMR (150 MHz, $CDCl_3$) δ 218.81, 193.72, 157.07, 142.60, 135.31, 126.74, 96.06, 71.99, 53.73, 50.81, 50.06, 47.88, 47.16, 42.27, 42.22, 42.02, 38.41, 35.33, 30.93, 29.57, 25.96, 22.73, 18.84, 18.22, 16.22; IR (thin film): 2357, 1738, 1664cm$^{-1}$; HRMS (ESI+) *m/z* calcd for $C_{25}H_{36}O_3Na$ $[M+Na]^+$: 407.2557, found: 407.2563.

[1]Ophiobolin I was kindly provided by Prof. Antonio Evidente (University of Naples, Italy).

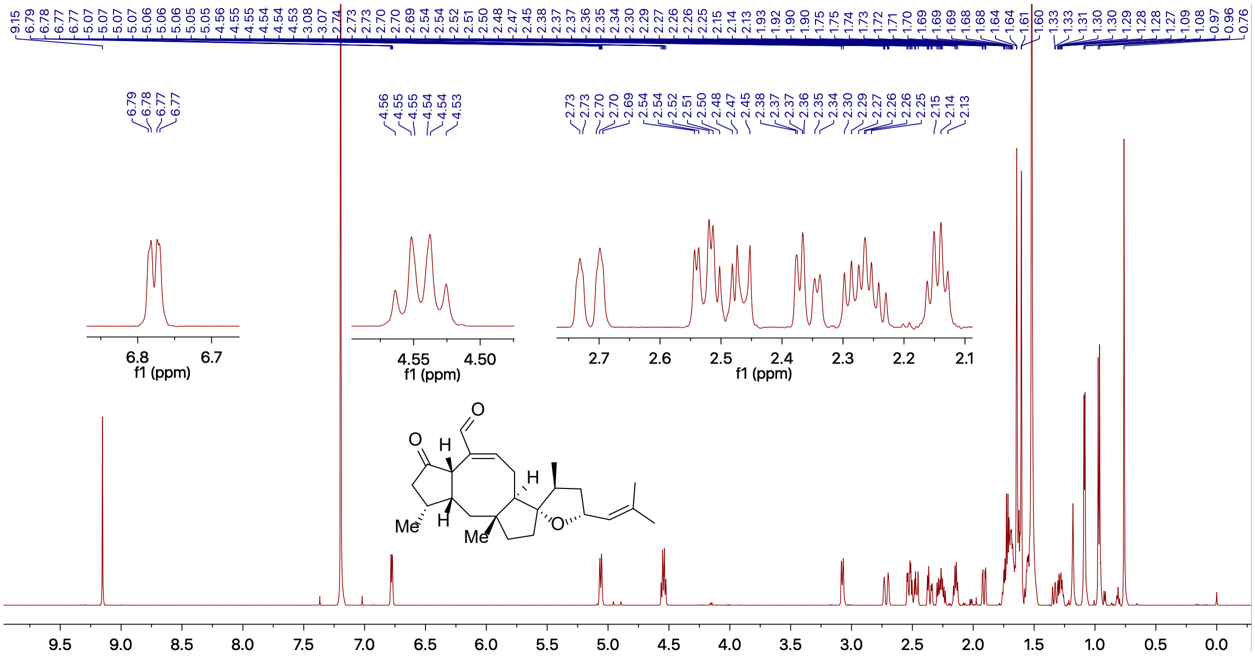


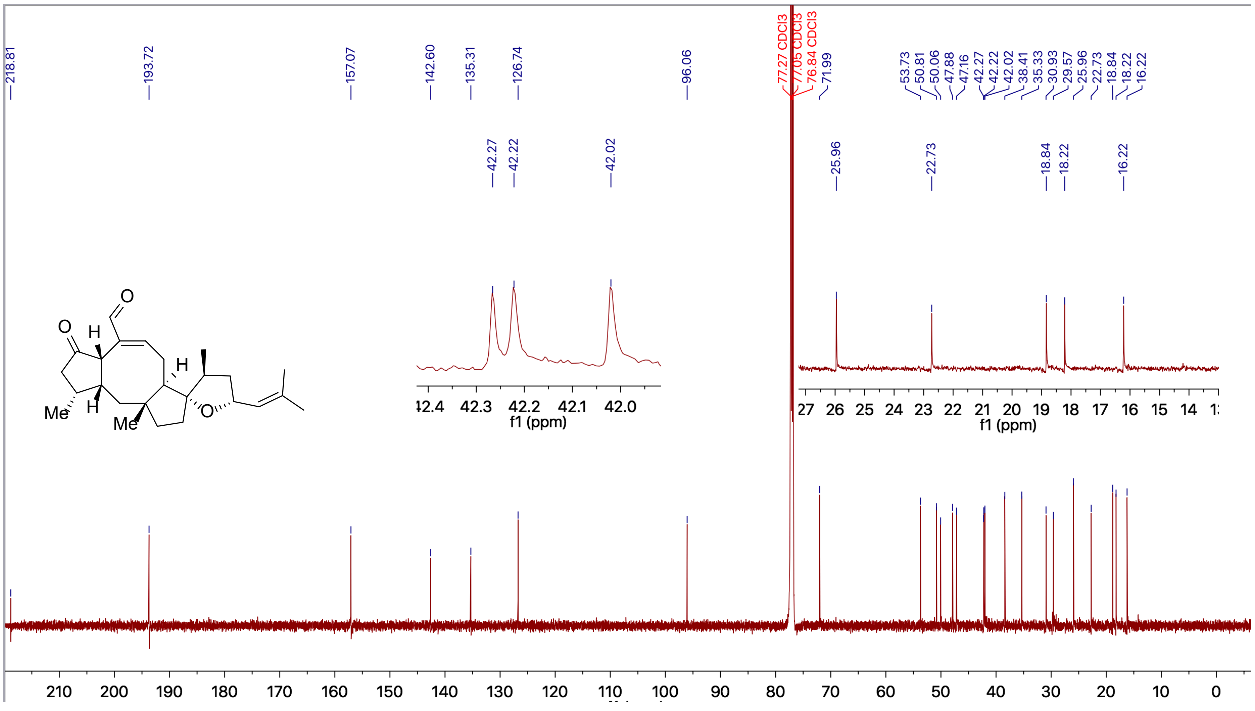


$^{1}H$ NMR (600 MHz, $CDCl_3$) and $^{13}C$ NMR (150 MHz, $CDCl_3$) of **3-deoxy-OpA**

Supporting Information

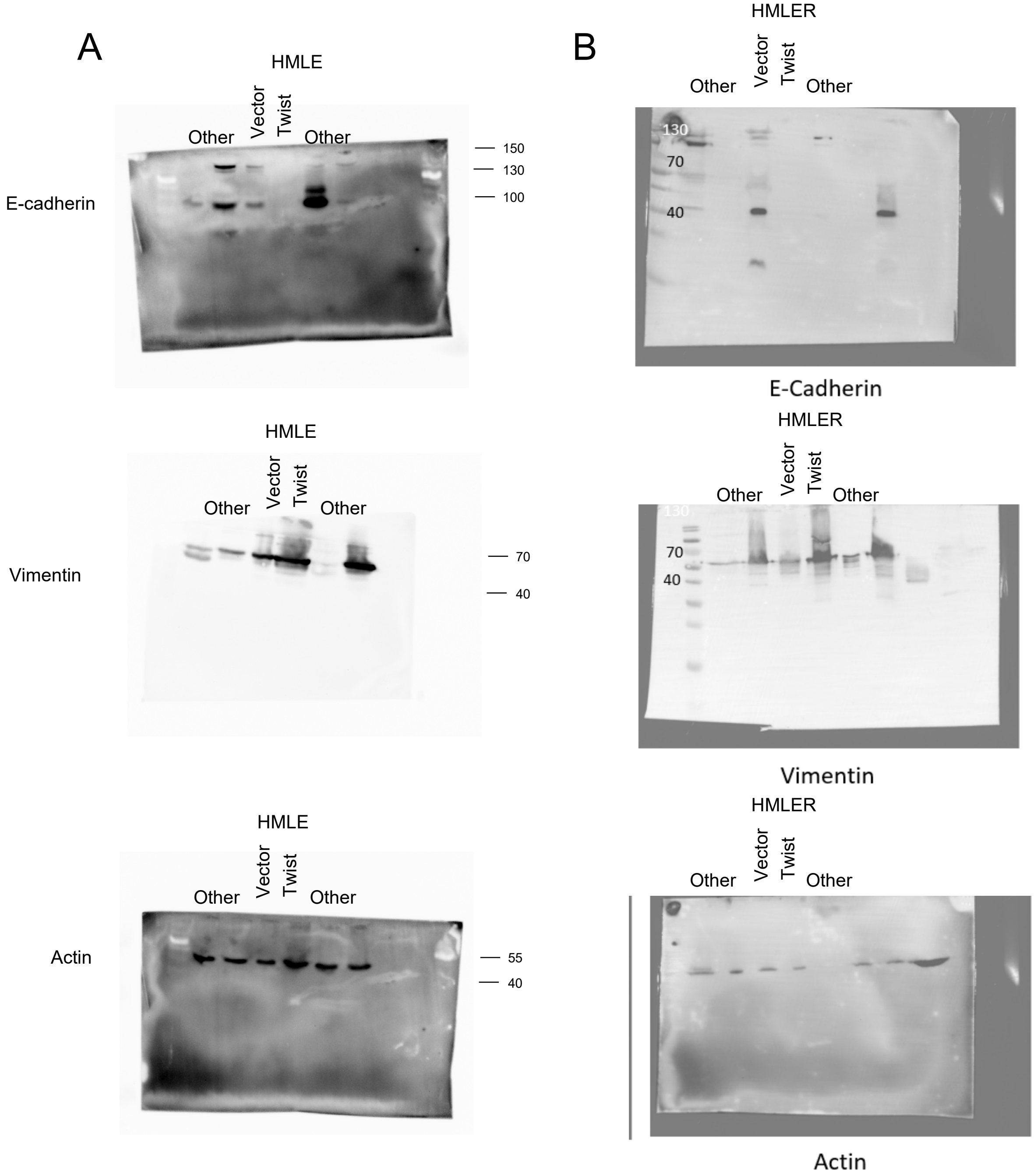


**Twist-induced protein expression.** Western blotting for the indicated proteins was performed on whole cell lysates from vector or Twist-transduced HMLE (A) or HMLER (B) cells. Chemiluminescence was detected and imaged. Molecular weight of ladder proteins, indicated on the right, was used to trim the membranes prior to horizontal membrane cutting for (B).
